# BNIP3 upregulation characterizes cancer cell subpopulation with increased fitness and proliferation

**DOI:** 10.1101/2022.03.21.485237

**Authors:** Yanyan Zhu, Bowang Chen, Junya Yan, Wendi Zhao, Pengli Dou, Na Sun, Yaokai Wang, Xiaoyun Huang

## Abstract

BNIP3 is a BH3 only protein with both pro-apoptotic and pro-survival roles depending on the cellular context. It remains unclear how BNIP3 RNA level dictates cell fate decisions of cancer cells. Here we undertook a quantitative analysis of BNIP3 expression and functions in single cell datasets of various epithelial malignancies. Our results demonstrated that BNIP3 upregulation characterizes cancer cell subpopulations with increased fitness and proliferation. We further validated the upregulation of BNIP3 in liver cancer organoids compared with 2D culture. Taken together, the combination of *in silico* perturbations using public single cell datasets and experimental cancer modeling using organoids ushered in a new approach to address cancer heterogeneity.

## Introduction

Heterogeneity of cancer is a well-known phenomenon that poses a daunting challenge for effective treatment. Cell-to-cell variability in signaling pathways and transcription factor activities renders the whole cancer population only partially responsive to most drugs[1, 2]. Design of better combination targeting strategy relies on accurate identification of key genes and pathways that defines cancer cell subpopulations with increased cancer hallmarks.

The ability of cancer cells to elicit uncontrolled proliferation and evade apoptosis requires a healthy mitochondrial network maintained through coordinated fission and mitophagy[3]. BNIP3 is involved in cellular responses to a multitude of different stresses through either apoptotic or non-apoptotic cell death[4]. BNIP3 also serves as an autophagy receptor that plays crucial roles in the removal of damaged mitochondria via interaction with ATG8. We have previously shown that phosphorylation of S17 and S24 in the LC3 interacting domains dictates whether BNIP3 signals apoptosis or pro-survival mitophagy[5]. However, it is still unclear how the RNA expression level of BNIP3 dictates cell fate decisions of cancer cells at single cell level.

Single cell RNA sequencing (scRNA-seq) has been harnessed to gain important insights into cancer heterogeneity and resulted in overwhelmingly rich datasets[6]. Almost all solid tumors and hematological malignancies have been investigated with scRNA-seq. Those datasets enabled the possibility to perform *in silico* perturbation experiments with single cell resolution to investigate the functional significance of genes of interest[7].

Here we undertook a comprehensive analysis of BNIP3 expression and functions in single cell datasets and TCGA datasets. We identified a cancer subpopulation characterized by upregulated BNIP3 in most epithelial malignancies. We also interrogated the pathway alterations in cancer cells with upregulated BNIP3 expression with a quantitative pathway enrichment approach using GSVA[8]. Our study underscored the power to combine computational and experimental approach to address gene-centered cancer heterogeneity.

## Results

BNIP3 expression was first investigated in the tumor and normal samples from the TCGA and the GTEx projects. Using transcripts per million reads normalization, BNIP3 expression was investigated in cancer samples and paired normal samples across different cancer types (**Supplementary Figure 1A**). The highest BNIP3 expression was found in KIRC, while significant patient-to-patient variability in BNIP3 was noted. Those population averaged measurements were incapable to capture the intratumoral heterogeneity reflected by cell-to-cell variability of cancer cells and heterogenous tumor ecosystem. Single cell transcriptomic datasets were used to determine the heterogeneous BNIP3 expression in cancer cells. Due to the inherent technical constrains of scRNA-seq, dropouts (zero UMI detected) were common. Considering the technical dropouts, cancer cells were stratified based on whether at least one UMI is detected whenever UMI count datasets were available. BNIP3 positivity actually might reflect BNIP3 upregulation. In almost all patients, scRNA-seq data revealed a cancer cell subpopulation with BNIP3 positivity.

The survival analysis was performed with all cancer types in the TCGA project (**Figure 1A**), suggesting BNIP3 mRNA expression as a worse prognostic factor also for Cervical Squamous Cell Carcinoma and Endocervical Adenocarcinoma (CESC), Cholangiocarcinoma (CHOL), Sarcoma (SARC). However, BNIP3 upregulation appeared to be a better prognosis indicator in Kidney Renal Clear Cell Carcinoma (KIRC) and Low Grade Glioma (LGG).

**Figure 1.**
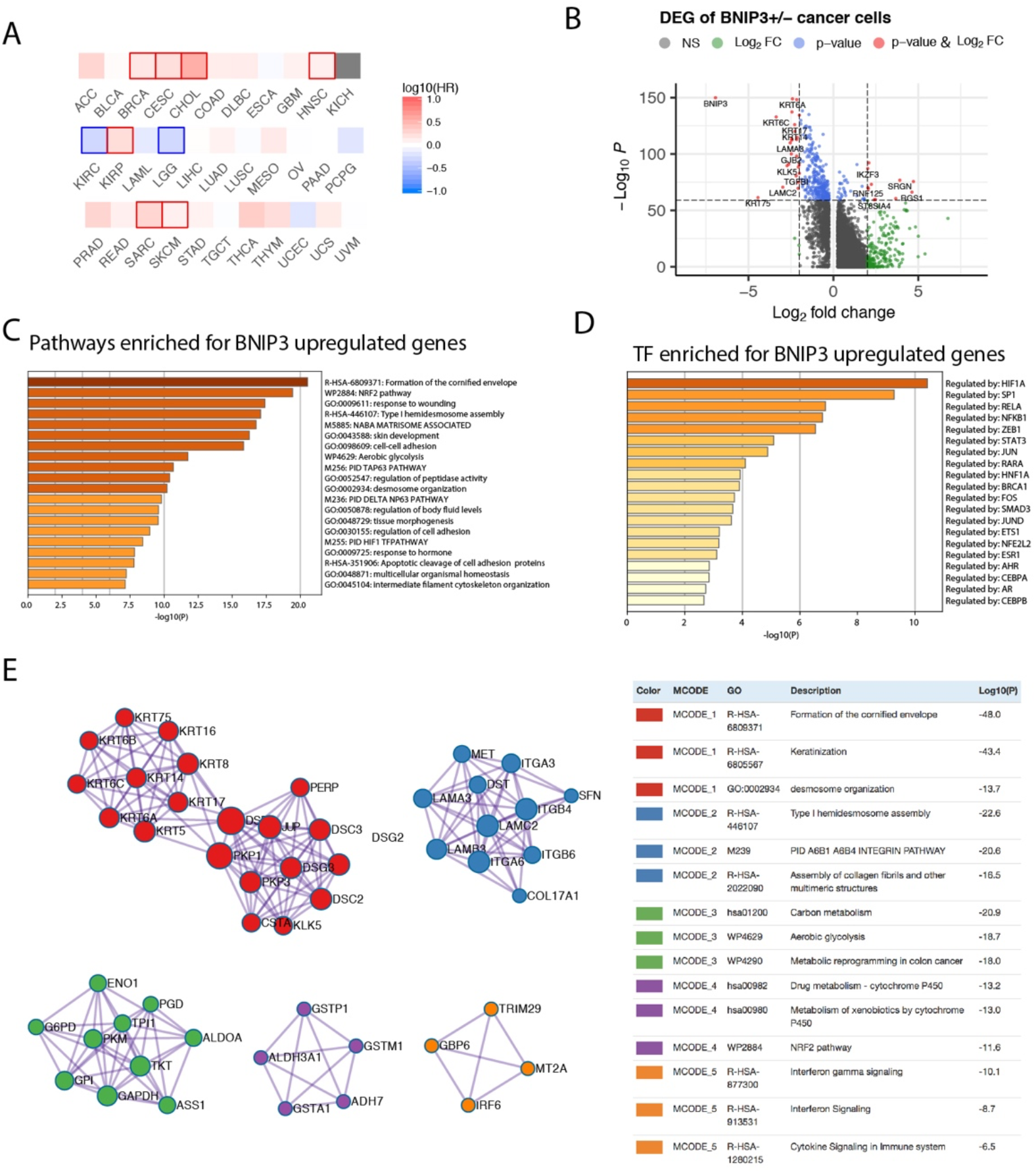
(A) Prognostic significance of BNIP3 in the TCGA cohort. Highlighted squares indicate p value smaller than 0.1. (B) Volcano plot showing the differentially expressed genes between BNIP3 positive and negative cancer cells. (C) Top pathways enriched for BNIP3 upregulated genes shown as barplot. (D) The top transcription factors enriched for BNIP3 upregulated genes. (E) Top protein-protein interaction modules enriched for BNIP3 upregulated genes.

The functional significance of BNIP3 in cancer cells was first investigated using a single cell dataset derived from head and neck cancer[9]. Cancer cells were stratified by BNIP3 RNA expression. The differentially expressed genes between BNIP3 positive and BNIP3 negative cancer cells were shown (**Figure 1B)**. The top pathways enriched for BNIP3 upregulated genes included formation of the cornified envelope, NRF2 pathway and response to wounding (**Figure 1C**). NRF2 is a transcription factor associated with antioxidant responses in cells. Interestingly, the top transcription factor regulating BNIP3 upregulated was HIF1A (**Figure 1D**), in agreement with the involvement of BNIP3 in cellular response to hypoxia. Using BNIP3 upregulated genes, protein-protein interaction network was constructed and analyzed for core modules. NRF2 pathway and metabolic reprogramming were among the enriched core modules, suggesting cancer cells with higher expression of BNIP3 might have achieved increased fitness by multiple pathways (**Figure 1E**).

To gain a quantitative insight into BNIP3 associated pathways, we employed gene set variation analysis to investigate the differential pathway activity of BNIP3 positive and negative cancer cells (**Figure 2A**). BNIP3 positive cancer cells have an upregulated activity in reactive oxygen species pathway, oxidative phosphorylation, as well as MYC targets. The high ROS burden within BNIP3 positive cancer cells might explain the feedback activation of antioxidant transcription factor NRF2. Cell cycle phase at the single cell level was inferred using single cell RNA, suggesting that the percentage of cells in S phase is higher in BNIP3 positive cells (**Figure 2B**). BNIP3 positive cancer cell subpopulation was also detected in lung cancer (**Supplementary Figure 1B**). Next, a lung cancer dataset with 42 patients was integrated with CCA or harmony algorithm and employed to obtain BNIP3 altered gene lists and pathways. Interestingly, BNIP3 upregulated genes were enriched for response to hypoxia, response to oxygen levels, response to decreased oxygen levels, and response to oxidative stress after CCA integration (**Figure 2C**). Response to oxidative stress was also among the top pathway enriched after harmony integration (**Figure 2D**).

**Figure 2.**
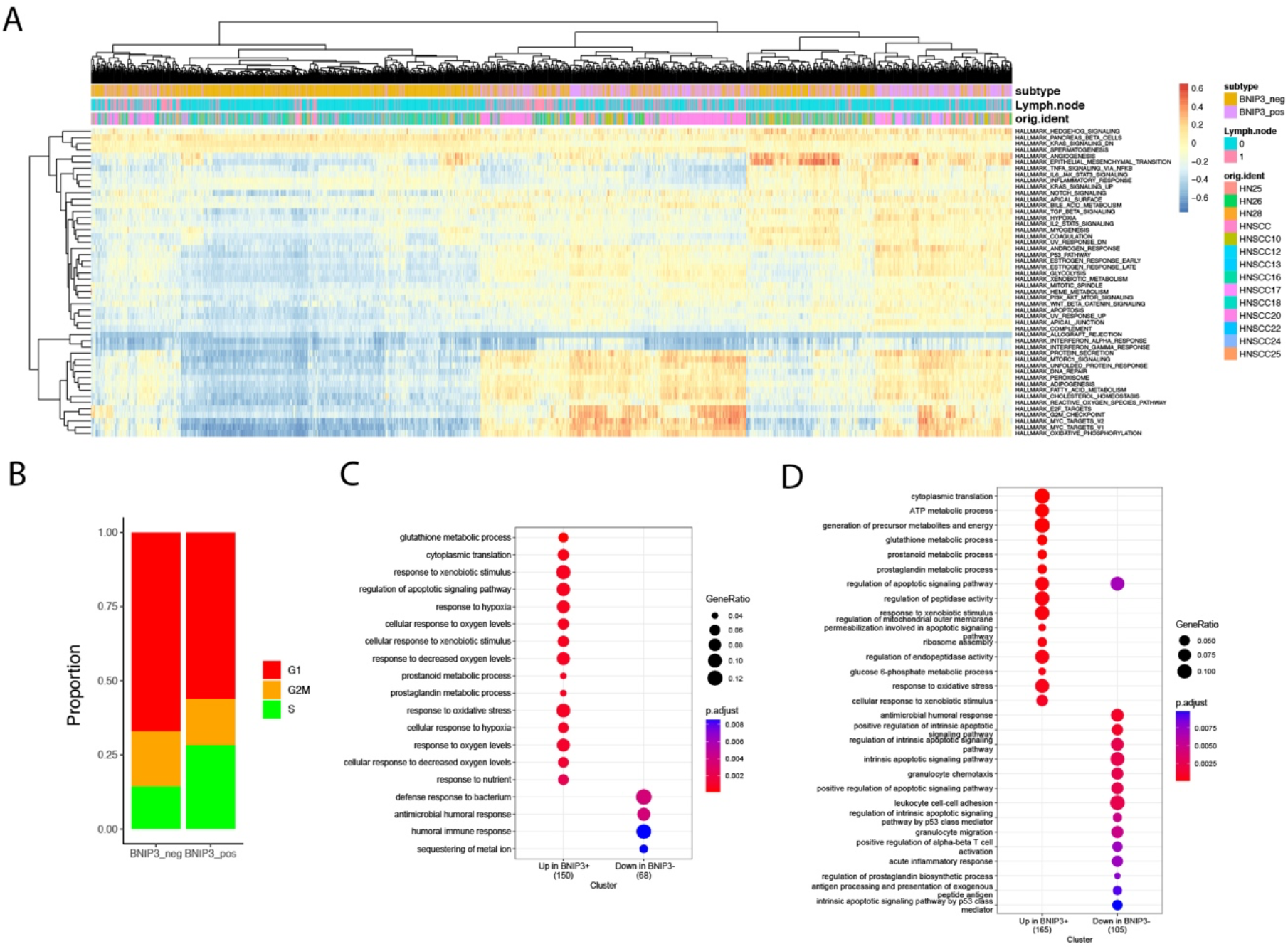
(A) Heatmap of the hallmark pathways at the single cell level. Each row represents one pathway and each column represents one cell. (B) Distribution of cell cycle phases for BNIP3 positive and BNIP3 negative cancer cells. (C) Pathway enrichment for BNIP3 up-regulated and down-regulated genes in lung cancer, using CCA integration. (D) Pathway enrichment for BNIP3 up-regulated and down-regulated genes in lung cancer, using harmony integration.

Next, we investigated BNIP3 expression in a cervical cancer single cell atlas. BNIP3 positive cervical cancer cells displayed a shifted transcriptional signature (**Figure 3A**). Cervical cancer patients with high BNIP3 expression in the TCGA cohort had a significantly decreased overall survival as compared with those with low BNIP3 expression(**Figure 3B**). The top 3 pathways enriched for BNIP3 upregulated genes were HIF1 TF pathway, response to wounding and Vitamin D receptor pathway (**Figure 3C**). The top 3 transcription factors regulating the upregulated genes were HIF1A, SP1 and RELA (**Figure 3D**). The proportion of BNIP3 positive cells is higher in breast cancer cells as compared to normal breast epithelial cells (**Figure 3E**). HER2 positive and triple negative breast cancers seemed to have an increased proportion of BNIP3 positive cancer cells as compared with ER positive breast cancers (**Supplementary Figure 1C**). Of note, BNIP3 is mostly expressed by epithelial cells, but not immune cells in the tumor microenvironment (**Figure 3F**). The prognostic significance of BNIP3 in breast cancer patients was also investigated in the TCGA breast cancer cohort. Patients with high BNIP3 expression had a significantly worse prognosis compared with patients with low BNIP3 expression (**Figure 3G**).

**Figure 3.**
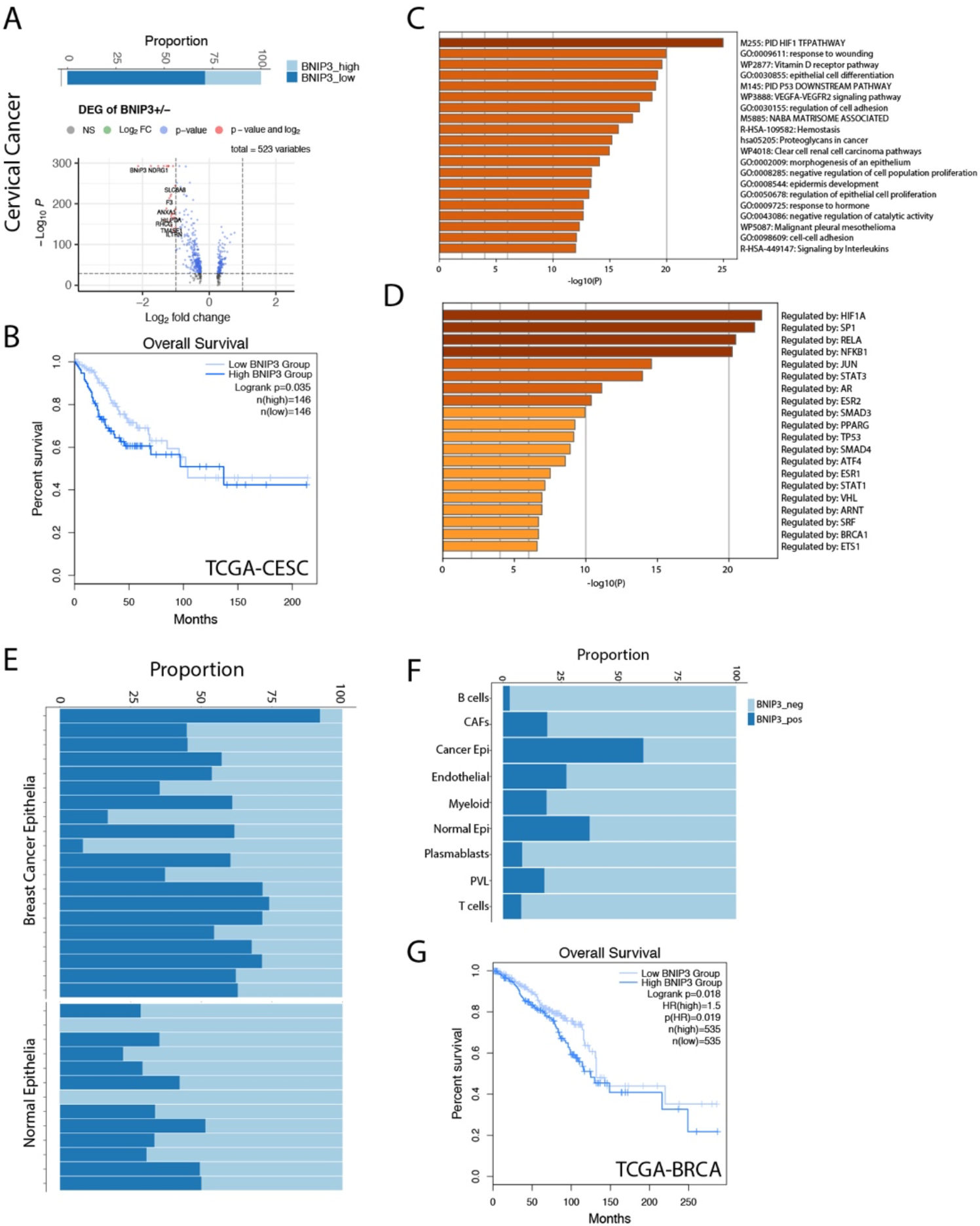
(A) The percentage of BNIP3 high and low cancer cells in cervical cancer and the volcano plot visualizing the differentially expressed genes between BNIP3 high and low cervical cancer cells. (B) Survival curve for cervical cancer patients in the TCGA cohort, stratified by BNIP3 mRNA expression. (C) Top pathways enriched for BNIP3 upregulated genes shown as barplot. (D) The top transcription factors enriched for BNIP3 upregulated genes. (E) Proportion of BNIP3 positive and negative cells in cancer epithelia and normal epithelia of the breast shown visualized as stacked barplot. Each bar indicates one individual. (F) Proportion of BNIP3 positive cells and negative cells in major cell types in the breast cancer cell atlas. (G) Survival curve for breast cancer patients in the TCGA cohort, stratified by BNIP3 mRNA expression.

The expression of BNIP3 was also investigated in a normal liver cell atlas. BNIP3 was mostly expressed by hepatocytes in the liver, but not by immune cells or stromal cells (**Figure 4A, 4B**). BNIP3 positive hepatocytes appeared to have more active cycling feature, as evidenced by an increased proportion of hepatocytes in S and G2M phase (**Figure 4C**). In the TCGA liver cancer cohort, we did observe that liver cancer patients with high expression of HIF1A or NRF2 (NFE2L2) tend to have worse prognosis (p value < 0.1) (**Figure 4D, 4E**). The expression of HIF1A and NRF2 was highly correlated in liver cancer samples from the TCGA cohort (**Figure 4F**).

**Figure 4.**
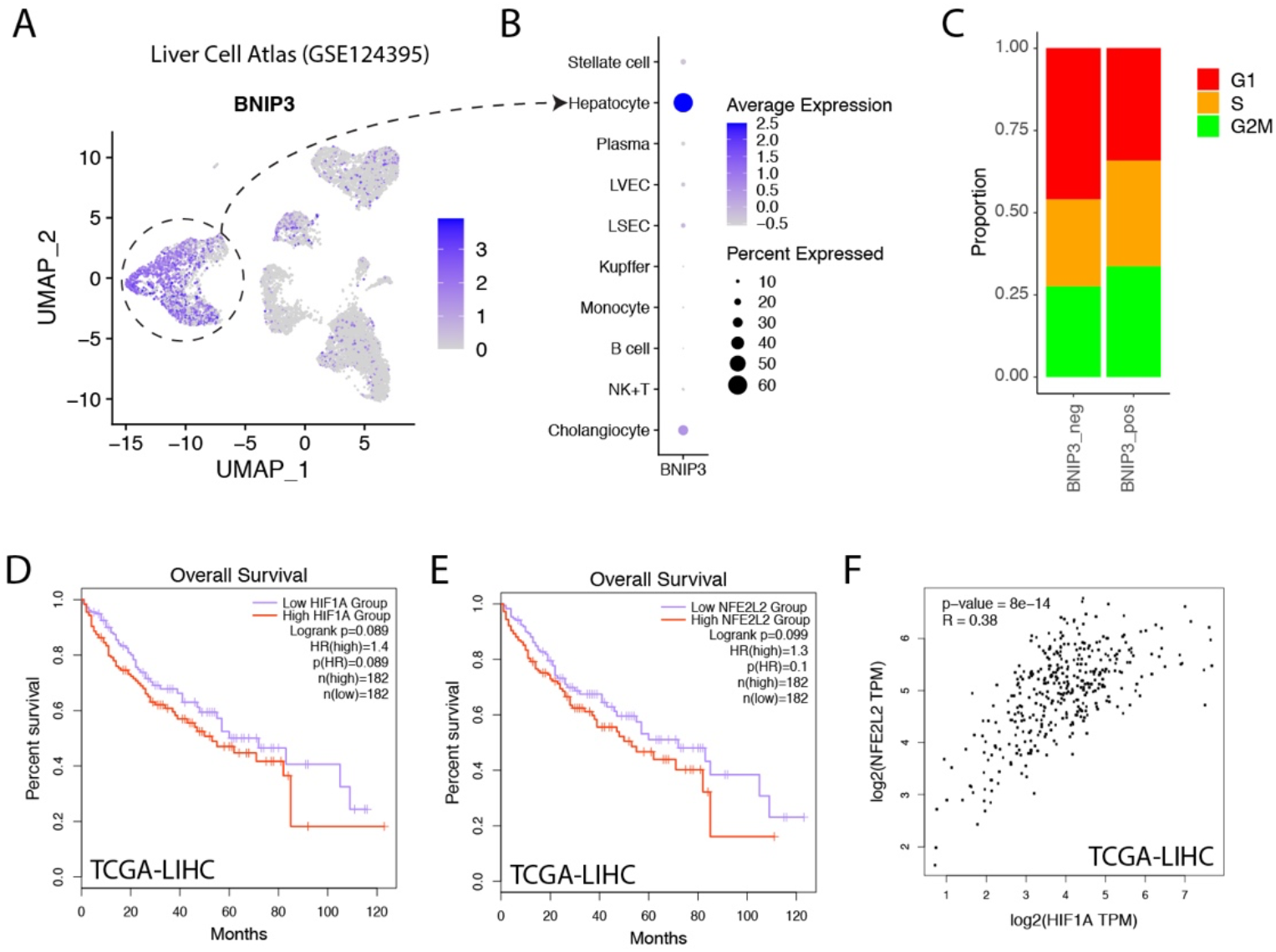
(A) Liver cell atlas visualized in UMAP plot, the intensity of color indicating expression of BNIP3. (B) Dotplot visualization of BNIP3 in major cell types within the liver. (C) Distribution of cell cycle phases for BNIP3 positive and BNIP3 negative hepatocytes. (D) Survival analysis of hepatocellular carcinoma patients in the TCGA cohort, stratified by mRNA expression of HIF1A. (E) Survival analysis of hepatocellular carcinoma patients in the TCGA cohort, stratified by mRNA expression of NFE2L2. (F) Correlation between the expression of HIF1A and NFE2L2 in liver cancers in the TCGA cohort.

Cancer cells cultured as organoids could better represent cancer cells grown in vivo and were shown to harbor increased stemness compared with cancer cells in 2D culture. We hypothesized that cancer cells might upregulate BNIP3 as a means to increase fitness when monolayer cell lines were converted into organoid lines. To validate this hypothesis, Hep G2 cell line was used as parental cell line to establish a liver cancer organoid line (**Figure 5A**). Hep G2 2D culture and organoid culture were subjected to bulk RNA-seq. The similarity matrix derived from RNA-seq data indicated a global change of transcriptome from 2D culture to 3D organoid culture (**Figure 5B**). Both oxidative phosphorylation and reactive oxygen species pathway increased in Hep G2 organoids compared with Hep G2 2D culture (**Figure 5C**). Liver cancer cells cultured as organoids have an upregulated CD24 expression, which played important roles in evasion from phagocytosis of cancer cells from macrophages (**Figure 5D**).

**Figure 5.**
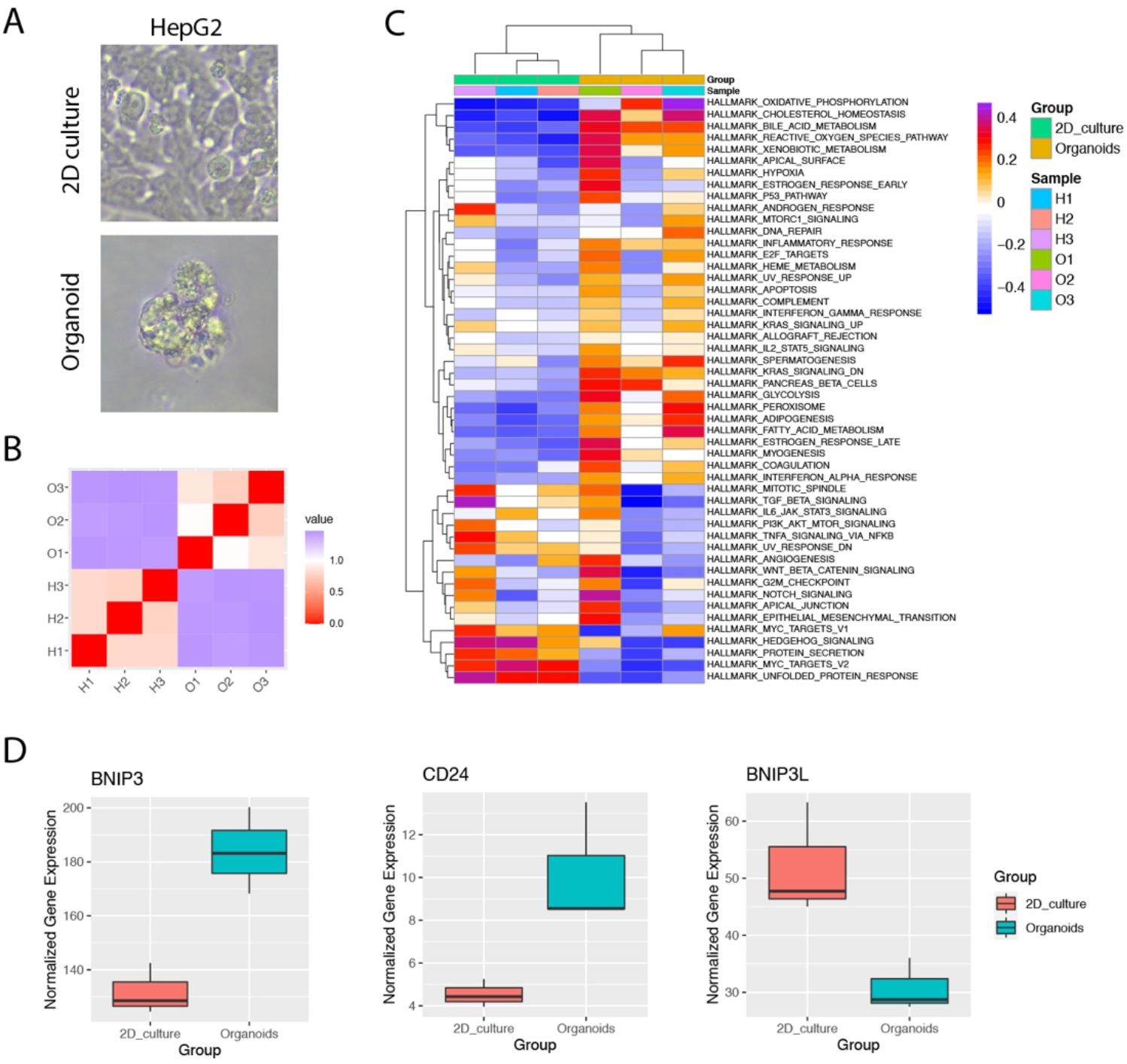
(A) Images of Hep G2 cancer cells cultured in 2D culture or organoid culture. (B) Heatmap of the correlation matrix between individual cancer transcriptomes derived form 2D culture or organoid culture. (C) GSVA analysis of hallmark pathways for individual cancer samples. Each row represents one hallmark pathway and each column represents one sample. Both rows and columns were arranged by hierarchical clustering. (D) Boxplots showing the expression of indicated genes for Hep G2 cultured in 2D or organoids.

## Discussion

As carcinogenesis progresses, cancer cells are making important decisions of life and death constantly. Cancer cells have unlocked the secret of phenotypic plasticity represented by distinct subpopulations with genetic or epigenetic variability. Identification of key genes and pathways that serve as master regulators of cancer cell fate decisions is key for the design of optimal treatment strategy. Our study unraveled BNIP3 upregulation as a hallmark characterizing cancer cell subpopulation with increased fitness and proliferation.

Single cell RNA sequencing has been applied by research community to gain insights into cancer heterogeneity and cellular ecosystem. The enormous datasets generated so far would serve as a gold mine to identify key regulators of cancer cell fate decisions if carefully reanalyzed and integrated.

Interestingly, the cancer type with the highest BNIP3 expression is clear cell renal cell carcinoma (ccRCC). This is in agreement that HIF is no longer degradable due to the loss of tumor suppressor VHL in ccRCC[10]. It has been demonstrated in vitro that siRNA mediated down-regulation of BNIP3 very effectively reduced the colony forming capacity of RCC cells[11]. BNIP3 overexpression has also been shown to enhance tumor growth for lung cancer[12] and liver cancer[13]. In liver cancer cells, BNIP3 was proposed to be a therapeutic target for cancer metastasis as BNIP3 upregulation enhanced anoikis resistance of HCC cells.

Cancer cells have harnessed the built-in cellular programs to adapt to hypoxia, which is a common feature of tumor microenvironment. The hypoxic niches typically renders chemotherapy[14] or radiation therapy[15] ineffective. Targeting HIF-2a with belzutifan (MK-6482) has been quite successful in a recent phase II trial, achieving a 49% objective response rate in patients with renal cell carcinoma[16]. Another key transcription factor NRF2 underlying BNIP3 upregulated cancer cell subpopulation has also recently been indirectly targeted with a chemical proteomic approach[17].

Our studied suggested BNIP3 might be involved in the enhanced tumorigenicity of liver cancer cells. This is consistent with a previous report that BNIP3 protects HepG2 cells against etoposide induced cell death under hypoxia[18]. Furthermore, BNIP3 upregulated cancer cells might be armed with immune evasion arsenals. Our results have demonstrated that CD24, a “don’t eat me” signal, has been upregulated in liver cancer organoids together with BNIP3. It has also been shown that hypoxia inducible factor elevated the expression of PD-L1 in ccRCC cells[19].

Taken together, the systems biology approach marrying *in silico* perturbations using public single cell datasets and experimental cancer modeling using organoids in our study unraveled a cancer cell subpopulation characterized by BNIP3 upregulation and revealed the potential druggable master regulators of enhanced fitness and proliferation.

## Methods

### Processing of Single Cell Datasets

For single cell datasets, annotations (meta data) from the original publications were used whenever possible. For GSE131907, “Malignant cells” as defined by original researchers were considered as cancer cells and used in our analysis. For GSE168652, cells with number of detected genes (nFeature_RNA) between 500 and 7500 were retained. The upper limit of total UMI count was set as 50000 to remove potential doublets and multiplets. Cells with more than 20% of mitochondrial RNA detected were also removed from our analysis. For datasets without meta data, quality control and unsupervised clustering was performed with Seurat. The count data was normalized using “LogNormalize” method with scaling factor of 10000. The top 2000 most variable genes were selected using “vst” method. For cancer cell grouping based on BNIP3 expression, cancer cells with at least one UMI detected for BNIP3 were considered as BNIP3 positive.

### Identification of Differential Expressed Genes

Upregulated genes in each cell cluster were identified using the FindMarkers function with statistical test method “wilcox”. Only genes expressed in more than 25% of cells and altered with log2FC higher than 0.25 were retained for further analysis.

### Inference of Cell Cycle Phase from Single Cell Data

Cell cycle scoring with single cell transcriptomic data was performed with CellCycleScoring function in Seurat. Each cell is assigned a score based on expression of G2/M markers and S phase markers. Cell cycle phase was predicted based on the respective cell cycles scores (G1, S, G2M). The genes used for cell cycle scoring is cc.genes.updated.2019 originally derived from a melanoma study[20].

### Gene list analysis

Differentially expressed genes with |log2FC| higher than 1 and p value smaller than 0.05 were subjected to gene list analysis using metascape[21], including pathway enrichment, analysis of protein-protein interaction and inference of transcription factors. Default parameters were used for implementation.

### Gene Set Variation Analysis

GSVA analysis was implemented with GSVA package in R. The hallmark pathways and KEGG pathways were retrieved from MSigDB. For transcript per million reads (TPM) expression data, “Gaussian” was used as the kernel for the non-parametric estimation of the cumulative distribution function of expression levels. For single cell datasets, the normalized data slot from RNA assay was used as input for GSVA analysis implemented also using “Gaussian” as the kernel for the non-parametric estimation of the cumulative distribution function of expression levels.

### SCENIC Analysis

SCENIC[22] was implemented with pySCENIC software. Transcription factors and corresponding target genes (regulon) were inferred based on co-expression of genes across cells. In brief, SCENIC infers TFs and their target genes from correlations between the expression of genes across cells. A TF and its target genes are defined as a regulon. The regulons are then refined by pruning targets based on enriched motifs. Finally, the activity of a regulon is measured by an AUCell value in each single cell. A high AUCell value indicates high activity and enrichment of a regulon in a cell.

### Transcription Factor Scoring

The bulk RNA-seq data from HepG2 2D culture and organoid culture was analyzed by a method previously developed for global transcription factor activity scoring[23]. For each transcription factor, the target genes with known regulation modes were extracted from TTRUST database[24], resulting in a list of genes activated by the transcription factor and a list of genes repressed by the transcription factor. The ratio between the median expression level of activated target gene and the median expression level of repressed target gene was calculated for each transcription factor and log2 transformed to obtain a final transcription factor score.

### TCGA/GTEx data mining

Investigation of BNIP3 expression in cancer samples and normal samples from TCGA or GTEx consortium was performed with GEPIA2 (http://gepia2.cancer-pku.cn/)[25]. For survival map analysis, the significance level of 0.05 was used and the median expression was used to stratify patients into high expression group and low expression group. In total, 33 different cancer types from the TCGA project were investigated.

### Cell Culture

Hep G2 cells were seeded in 10 cm culture dish and maintained in DMEM medium (L110KJ, BasalMedia) supplemented with 10% FBS. Medium was renewed every two days. For derivation of organoid line, Hep G2 cells were centrifuged at 500 g for 5 minutes at 4 degree. The cell pellet was resuspended in Matrigel (R&D, 3533-005-02). For one well of 24-well plate, 50 ul cell suspension with 10000 cells was seeded for the Matrigel to solidify. After Matrigel solidification, 1 ml medium was added to each well. The organoid medium A contained 1% PS, 1% Glutamax, 10 mM HEPES, B27 (1:50), N2 (1:100), 1.25 mM n-Acetyl-L-cysteine, 10 mM nicotinamide, 10 nM recombinant human Gastrin I, 50 ng/ml recombinant human EGF, 100 ng/ml recombinant human FGF10, 25 ng/ml recombinant human HGF, 10 uM Forskolin, 5 uM A8301, 10 uM Y27632, 3 nM Dexamethasone. The organoid medium B contained 1% PS, 1% Glutamax, 10 mM HEPES, B27 (1:50), N2 (1:100), 1.25 mM n-Acetyl-L-cysteine, 10% Rspo-1 supernatant, 10 mM nicotinamide, 10 nM recombinant human Gastrin I, 50 ng/ml recombinant human EGF, 100 ng/ml recombinant human FGF10, 25 ng/ml recombinant human HGF, 10 uM Forskolin, 5 uM A8301.

The reagents used in our cell culture is as following:

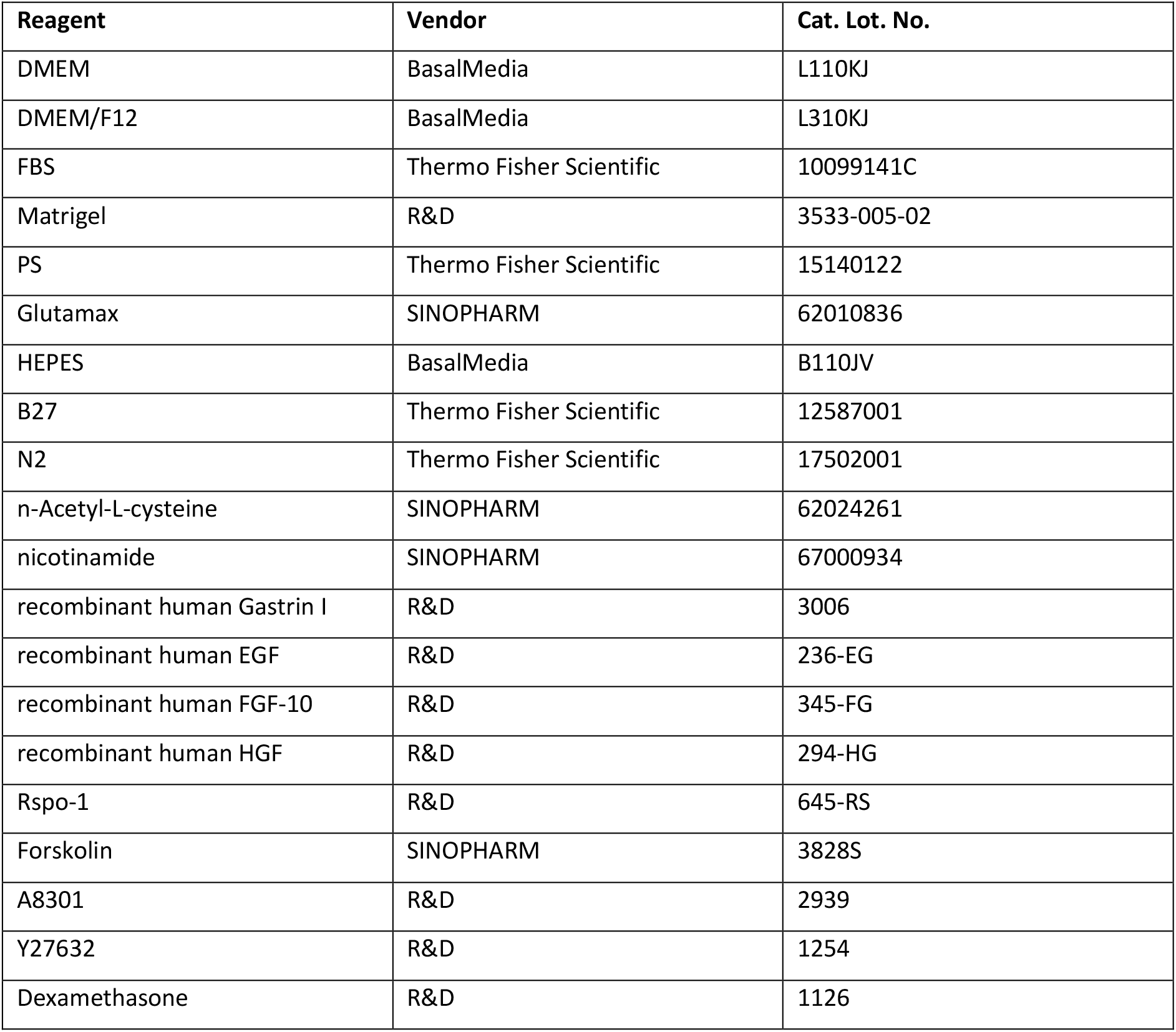

| Reagent | Vendor | Cat. Lot. No. |
| --- | --- | --- |
| DMEM | BasalMedia | L110KJ |
| DMEM/F12 | BasalMedia | L310KJ |
| FBS | Thermo Fisher Scientific | 10099141C |
| Matrigel | R&D | 3533-005-02 |
| PS | Thermo Fisher Scientific | 15140122 |
| Glutamax | SINOPHARM | 62010836 |
| HEPES | BasalMedia | B110JV |
| B27 | Thermo Fisher Scientific | 12587001 |
| N2 | Thermo Fisher Scientific | 17502001 |
| n-Acetyl-L-cysteine | SINOPHARM | 62024261 |
| nicotinamide | SINOPHARM | 67000934 |
| recombinant human Gastrin I | R&D | 3006 |
| recombinant human EGF | R&D | 236-EG |
| recombinant human FGF-10 | R&D | 345-FG |
| recombinant human HGF | R&D | 294-HG |
| Rspo-1 | R&D | 645-RS |
| Forskolin | SINOPHARM | 3828S |
| A8301 | R&D | 2939 |
| Y27632 | R&D | 1254 |
| Dexamethasone | R&D | 1126 |

### RNA Sequencing

After quality control with gel electrophoresis and Agilent 2100, mRNA were captured with beads coupled with oligo(dT) and fragmented before priming with random hexamers. First cDNA strand and second strand were synthesized and purified. The purified double stranded cDNA were subjected to end repairing, A-tailing and adapter ligation. The products were purified and size-selected before final PCR amplification. The PCR products were purified to obtain the final libraries, which were sequenced with Nova-seq 6000 to obtain 6G data for each sample. The raw reads were pre-processed and filtered before alignment to hg38 reference genome. Stringtie were employed to derive TPM expression matrix[26].

## Data availability

The single cell datasets analyzed in this study can be accessed from GEO database with the following accession numbers: head and neck cancer (**GSE103322**)[9], lung cancer (**GSE131907**)[27],lung cancer (**GSE148071**)[28], breast cancer (**GSE176078**)[29], cervical cancer (**GSE168652**)[30], normal liver (**GSE124395**)[31].

## Acknowledgement

We thank Yantao Du for providing the Hep G2 cell line used in our study. We also thank all the members of Center for Systems Biology (Intelliphecy) for discussion and input.

---

**Supplementary Figure 1.**
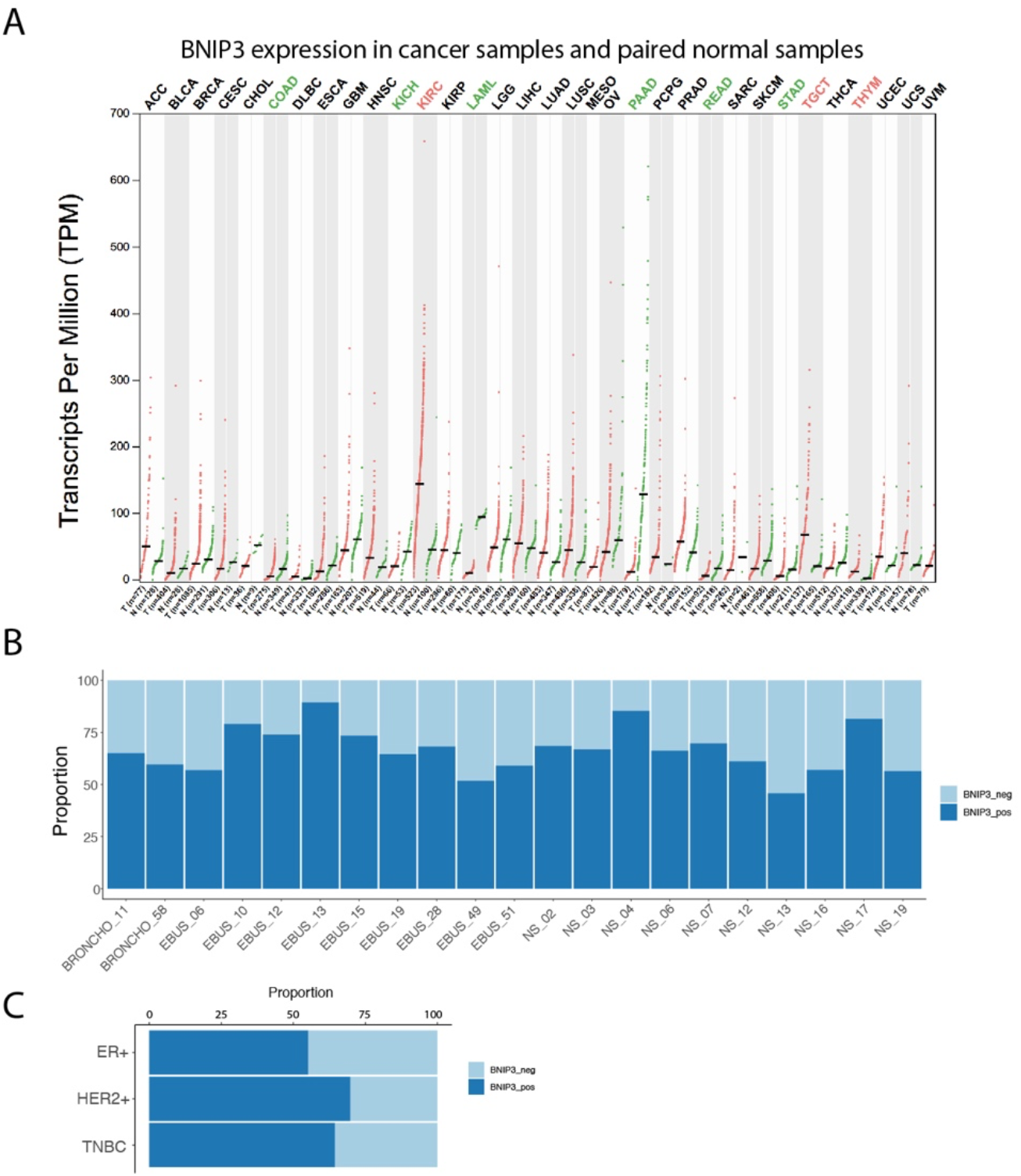

